# Selective inhibition of protein secretion by abrogating receptor-coat interactions during ER export

**DOI:** 10.1101/2022.01.31.478454

**Authors:** Natalia Gomez-Navarro, Julija Maldutyte, Kristina Poljak, Sew-Yeu Peak-Chew, Jonathon Orme, Brittany J. Bisnett, Caitlin H. Lamb, Michael Boyce, Davide Gianni, Elizabeth A. Miller

## Abstract

Protein secretion is an essential cellular process that permits cell growth, movement and communication. Traffic of proteins within the eukaryotic secretory pathway is mediated by transport intermediates that bud from one compartment, populated with appropriate cargo proteins, and fuse with a downstream compartment to deliver their contents. Here, we explore the possibility that protein secretion can be selectively inhibited by perturbing protein-protein interactions that drive capture into transport vesicles. Human PCSK9 is a major determinant of cholesterol metabolism, whose secretion is mediated by a specific cargo adaptor of the ER export machinery, SEC24A. We map a series of protein-protein interactions between PCSK9, its ER export receptor, SURF4, and SEC24A, that mediate secretion of PCSK9. We show that the interaction between SURF4 and SEC24A can be inhibited by 4-PBA, a small molecule that occludes a cargo-binding domain of SEC24. This inhibition reduces secretion of PCSK9 and additional SURF4 clients that we identify by mass spectrometry, leaving other secreted cargoes unaffected. We propose that selective small molecule inhibition of cargo recognition by SEC24 is a potential therapeutic intervention for atherosclerosis and other diseases that are modulated by secreted proteins.

## Introduction

Approximately a third of human protein-coding genes encode proteins that enter the secretory pathway. Such secretory proteins include integral membrane proteins, soluble proteins located in the lumen of membrane-bound organelles, and proteins that are transported outside of the cell. The vast majority of secretory proteins enter the pathway at the endoplasmic reticulum (ER), where they reach their mature state after acquiring post-translational modifications and interacting with the abundant chaperones of the ER lumen. Secretion-competent mature proteins are exported from the ER via vesicles generated by the COPII (coat protein complex II) coat and delivered to the Golgi apparatus, where they are further modified (Barlowe & Miller, 2013). Vesicle biogenesis occurs via self-assembly of the COPII coat at ER exit sites (ERES). Protein capture into COPII vesicles occurs via two mechanisms: signal-mediated export and bulk flow (Barlowe & Helenius, 2016). In signal-mediated export, specific signals within cargo proteins are recognized by the COPII coat, which then concentrates cargo at sites of vesicle formation. The coat-cargo interaction is direct for many transmembrane proteins, whereas soluble proteins or proteins that do not expose cytoplasmic signals require export receptors that bridge the interaction between cargo and coat (Geva & Schuldiner, 2014). In contrast, bulk flow export occurs in the absence of signals, and relies on diffusion of a cargo molecule into an ER exit site followed by passive uptake into a nascent vesicle (Thor *et al*, 2009; Wieland *et al*, 1987). The extent to which signal-mediated versus bulk flow mechanisms are used for the entire secretome remains to be fully determined.

The COPII coat subunit, Sec24, is the cargo adaptor that confers selective cargo capture and enrichment during vesicle formation. Mutagenesis and structural studies have revealed multiple cargo binding sites on Sec24 (Mancias & Goldberg, 2008; Mossessova *et al*, 2003; Miller *et al*, 2003), which permits recognition of diverse cargoes by a single coat subunit (Geva & Schuldiner, 2014). The repertoire of cargo molecules recognised by the coat is further expanded by multiple isoforms of Sec24: yeast express three Sec24 homologs (Sec24, Iss1/Sfb2 and Lst1/Sfb3); humans express four isoforms (Sec24A-D). Sec24A and Sec24B share 60% sequence identity, and Sec24C and Sec24D similarly share more than 50% sequence identity. Accordingly, the four isoforms can be classified into two subfamilies, Sec24A/B and Sec24C/D, that share non-overlapping recognition specificity (Tang *et al*, 1999; Pagano *et al*, 1999; Adolf *et al*, 2019; 2016).

The specificity of Sec24-mediated cargo recognition raises the possibility that secretion, an essential process, might be selectively targeted by preventing a subset of Sec24-cargo interactions. Indeed, release of the secreted proprotein convertase subtilisin kexin 9 (PCSK9) was selectively perturbed in SEC24A knock-out mice (Chen *et al*, 2013). Because PCSK9 participates in cholesterol homeostasis, the predominant physiological effect of SEC24A knock-out was reduced circulating cholesterol, suggesting a novel therapeutic avenue for cardiovascular health. Here, we sought to probe the network of protein-protein interactions that contribute to PCSK9 secretion with the goal of understanding what interactions might be perturbed to achieve a therapeutic outcome.

PCSK9 secretion via SEC24A is facilitated by the ER export receptor, SURF4, which also mediates secretion of multiple calcium-binding extracellular matrix proteins that employ N-terminal ϕ-P-ϕ signals, where ϕ is any hydrophobic amino acid (Emmer *et al*, 2018; Yin *et al*, 2018). This motif has been termed the ER-ESCAPE (<u>ER</u>-<u>E</u>xit by <u>S</u>oluble <u>C</u>argo using <u>A</u>mino-terminal <u>P</u>eptide-<u>E</u>ncoding) motif (Yin *et al*, 2018). The precise mechanism by which SURF4 recognises this signal remains unknown, and the nature of the interaction between SURF4 and SEC24A also remains unclear. Here, we demonstrate that the SEC24A-SURF4 interaction is driven by a conserved cargo-binding domain on SEC24A known as the B-site (Mossessova *et al*, 2003; Miller *et al*, 2003), which can be occluded by small molecule treatment (Ma *et al*, 2017) to selectively inhibit secretion of PCSK9 and additional SURF4 clients. Moreover, we identify new clients of SURF4 that, like PCSK9, use N-terminal ER-ESCAPE motifs to achieve SURF4-dependent secretion. The network of client-receptor-adaptor interactions revealed by our study lends support to the concept of selective inhibition of protein secretion as a therapeutic target.

## Results

### PCSK9 secretion requires both SEC24A and SURF4

Given the previously described roles for SEC24A and SURF4 in PCSK9 secretion (Chen *et al*, 2013; Emmer *et al*, 2018), we first evaluated the effects of deletion of these export factors on PCSK9 secretion in TREx-293 cells heterologously expressing V5-tagged (PCSK9-V5). We performed pulse-chase experiments to compare the kinetics of secretion in wild-type and knock-out cell lines. As expected, deletion of either SEC24A or SURF4 resulted in decreased release of PCSK9-V5 to the media, with no additive effect observed in the double knock-out (Figure 1A). Importantly, proteolytic processing of PCSK9-V5, which is required for secretion competence, remained unperturbed in the knock-out lines (Figure 1A, B). Secretion defects were rescued by re-introduction of SEC24A and SURF4 cDNAs by transient transfection (Figure S1A, B). We also tested PCSK9-V5 secretion in knock-out lines deleted for the remaining SEC24 isoforms, SEC24B, SEC24C, and SEC24D, finding that only SEC24A knock-out reduced PCSK9-V5 secretion (Figure S1C, D). These findings differ from previous results that used steady-state immunoblotting to test PCKS9 secretion in siRNA knockdown conditions, where depletion of SEC24B and SEC24C, also perturbed PCSK9 secretion (Deng *et al*, 2020).

**Figure 1:**
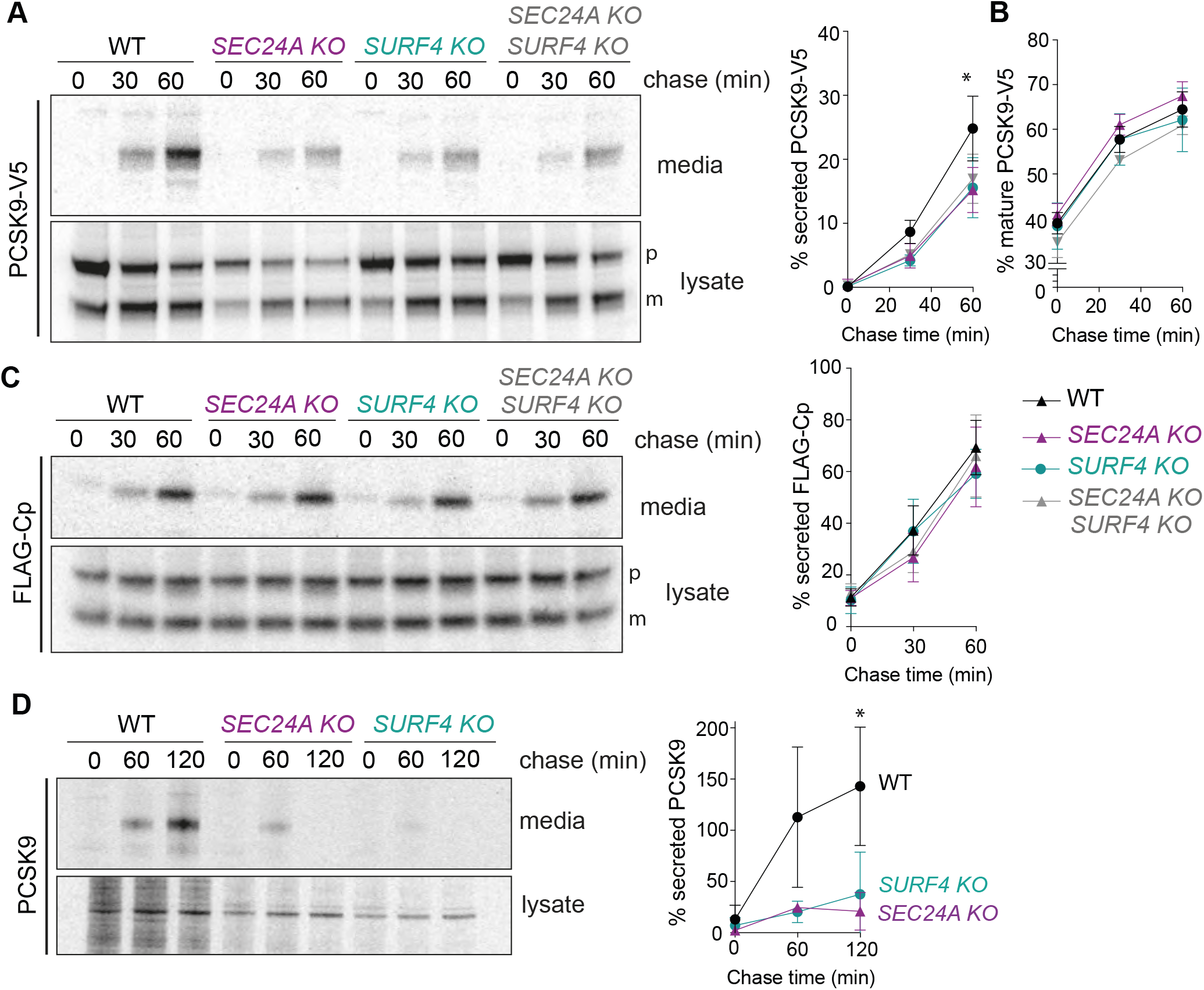
PCSK9 secretion requires SEC24A and SURF4. **(A)** PCSK9-V5 maturation and secretion was examined in WT and KO cell lines by pulse chase with [^35^S]methionine. PCSK9-V5 was immunoprecipitated with α-V5 from lysates and conditioned media at the indicated times and detected by SDS-PAGE and autoradiography. Percentage of secreted PCSK9 was calculated as [band intensity in media at a given time point]/[total protein intensity (media + lysate) at time 0]. Plots show the mean ± SD of three independent experiments. Statistical test was a one-way ANOVA with Dunnett’s correction for multiple comparisons, * p<0.05, ** p<0.01, *** p<0.001. **(B)** Percentage of mature PCSK9 was calculated as [band intensity of the lower MW (mature) species at a given time point]/ [total protein (mature + proprotein + secreted) at time 0]. Plots show the mean ± SD of three independent experiments. Statistical test was a one-way ANOVA with Dunnett’s correction for multiple comparisons, * p<0.05, ** p<0.01, *** p<0.001. **(C)** FLAG-Cp secretion was examined in WT and KO cell lines. FLAG-Cp was immunoprecipitated with α-FLAG from lysates and conditioned media at the indicated times and detected by SDS-PAGE and autoradiography. Autoradiographs are representative of three independent experiments. As in A, secretion of FLAG-Cp was quantified by phosphorimage analysis. Plots of the mean ± SD are shown in the right-hand panel. **(D)** Endogenous PCSK9 secretion from the indicated HuH7 cell lines was quantified by pulse chase with [^35^S]methionine. PCSK9 was immunoprecipitated with α-PCSK9 from lysates and conditioned media at the indicated times and detected by SDS-PAGE and autoradiography. Plots of the mean ± SD of three independent experiments are shown in the right-hand panel.

We measured global secretion using pulse-chase analysis of a FLAG-tagged bulk flow marker, the C-terminal protease domain (Cp) of the capsid protein of Semliki Forest virus (Thor *et al*, 2009). FLAG-Cp was secreted normally in both *SEC24AKO* and *SURF4KO* cell lines, suggesting bulk secretion remained intact (Figure 1C). Finally, knock-out of either SEC24A or SURF4 in the hepatic cell line HuH7 that endogenously expresses PCSK9, resulted in an even more dramatic reduction in PCSK9 secretion (Figure 1D). The magnitude of this effect suggests that in cells specialised for PCSK9 secretion, both SEC24A and SURF4 play essential functions in PCSK9 ER export.

### SURF4 interacts with the B-site of SEC24

In order to characterise the mechanisms of selective SEC24A-and SURF4-mediated secretion we sought to characterise the interaction between SURF4 and SEC24A. We employed a cell-based protein-protein interaction assay based on the NanoBiT (<u>Nano</u>Luc <u>Bi</u>nary <u>T</u>echnology) enzymatic complementation technology. We fused the 2 fragments of NanoLuc, Large BiT (LgBiT, 18 kDa) and Small BiT (SmBiT, 11 amino acids), to the N- and C-termini of SEC24A and SURF4, and assayed luminescence complementation of the paired tagged fusions (Figure 2A). The different linkage orientations revealed optimal signal with SmBiT appended to the N-terminus of SEC24A and LgBiT appended to the N-terminus of SURF4 (Figure 2A). Luminescence from the SEC24A-SURF4 pair was 4-fold higher than luminescence for a control interaction that paired LgBiT-SURF4 with cytoplasmic HaloTag-SmBiT control (Figure 2B). Tagging of either SEC24A or SURF4 at C-terminus resulted in lower levels of luciferase activity (Figure 2B, S2A). To demonstrate the specificity of the interaction, we measured the luminescent signal upon increasing expression of untagged SURF4, which indeed decreased luminescence, presumably by competing for interaction with SmBiT-SEC24A (Figure S2B).

**Figure 2:**
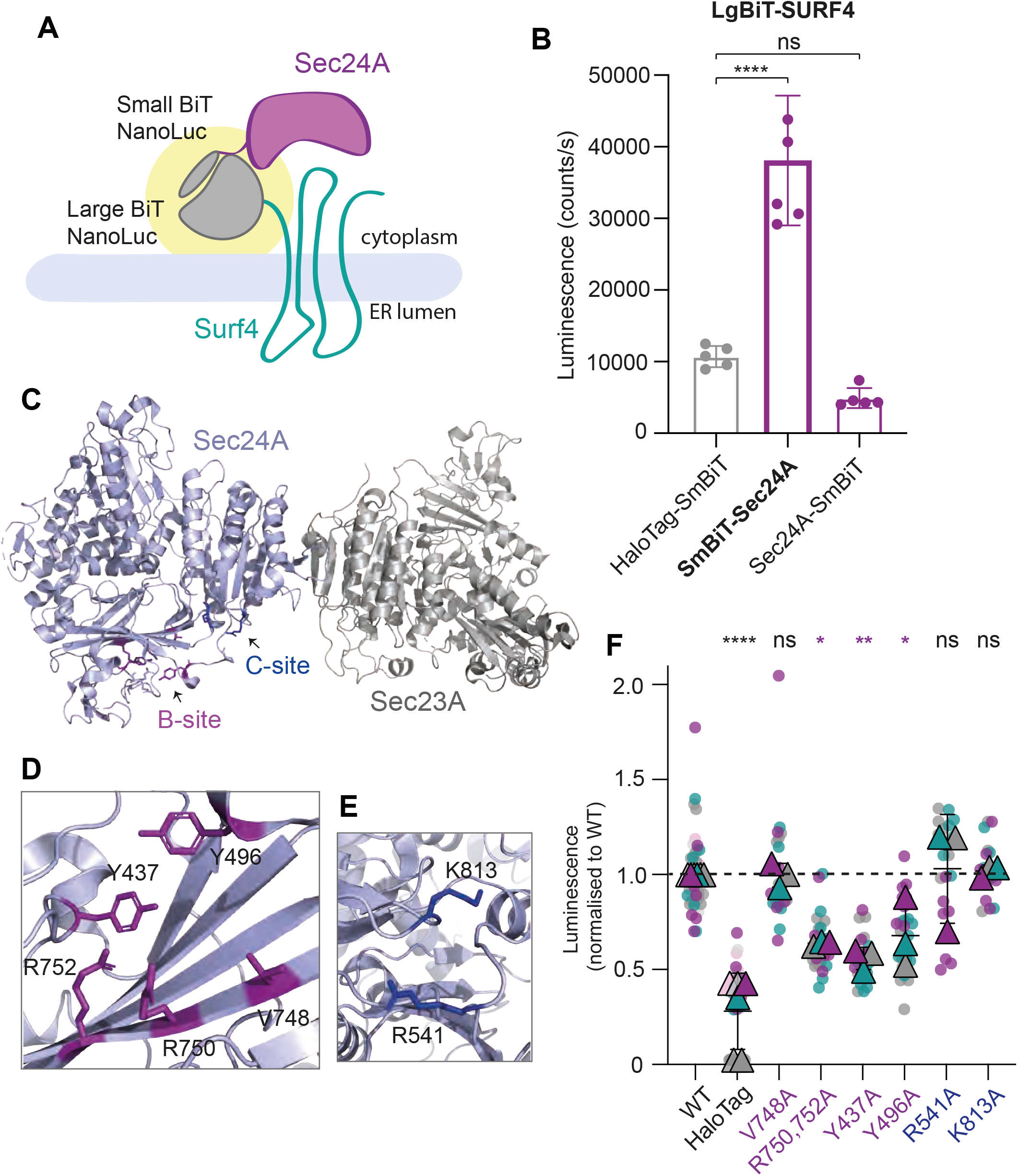
SURF4 and SEC24A interact via the B-site of Sec24. **(A)** Cartoon of the NanoBiT complementation assay for measuring SEC24A and SURF4 interaction, where complex formation results in a reconstituted luminescence signal (yellow halo). **(B)** Cellular luminescence was measured for the indicated NanoBiT constructs expressed in *SEC24A SURF4* KO cells. Graph shows the mean luminescence ± SD (n = 6); statistical test was a one-way ANOVA with Dunnett’s correction for multiple comparisons, * p<0.05, ** p<0.01, *** p<0.001, **** p<0.0001. **(C)** Crystal structure of the SEC24A-SEC23A complex (PDB 5VNO. Residues in the B-site are shown in purple and residues in the C-site are shown in blue. **(D, E)** Structural detail of the B and C sites, respectively, with key residues shown as sticks. **(F)** Luminescence values measured upon co-expression of LgBiT-SURF4 with the indicated SmBiT-SEC24A mutants normalised to WT values. Data correspond to six technical replicates per biological replicate, and three independent experiments. The plotted triangles represent the mean ± SD of the 3 biological replicates. Statistical test was a one-way ANOVA with Dunnett’s correction for multiple comparisons, * p<0.05, ** p<0.01, *** p<0.001, **** p<0.0001.

We next mutated surface-exposed residues on SEC24A previously described as cargo binding sites (Mancias & Goldberg, 2008; Mossessova *et al*, 2003; Miller *et al*, 2003) and screened for mutants that reduced the reconstituted luminescence signal of the SEC24A-SURF4 NanoBiT pair. We targeted two well-characterised cargo-interaction sites, the B- and C-sites (Figure 2C), making multiple mutations in each region, none of which perturbed protein stability (Figure 2D, E; Figure S2C). Mutations in the C-site (R541 and K813) did not perturb the interaction between SEC24A and SURF4, however, mutation of four residues that line a conserved pocket known as the B-site (R750, R752, Y437 and Y496) significantly reduced the luminescence signal, suggesting impaired interaction between SEC24A and SURF4 (Figure 2F).

### SURF4-SEC24A interaction is disrupted by 4-PBA

Having identified the SEC24A B-site as driving the interaction between SEC24A and SURF4, we tested whether occlusion of the B-site by 4-phenylbutyrate (4-PBA) (Ma *et al*, 2017) perturbs interaction with SURF4. Indeed, in our NanoBiT assay, increasing concentrations of 4-PBA resulted in a reduction in luminescence, suggesting that 4-PBA competes with SURF4 at the B-site (Figure 3A). 4-PBA at those concentrations did not affect cell viability (Figure S3A). We next asked whether impaired SEC24A-SURF4 interaction caused by 4-PBA resulted in deficient trafficking of SURF4-dependent cargo proteins. We performed pulse-chase experiments to measure PCSK9 secretion in the presence of 10 mM 4-PBA (Figure 3B). Indeed, 4-PBA significantly reduced PCSK9 secretion while not affecting protein maturation (Figure 3C), suggesting that PCSK9 secretion indeed relies on the B-site of Sec24 for its ER export. We also analysed the effects of 5-PVA, a 4-PBA analog that has a similar affinity for Sec24 11, and observed a comparable reduction in the SEC24A-SURF4 interaction (Figure S3B).

**Figure 3:**
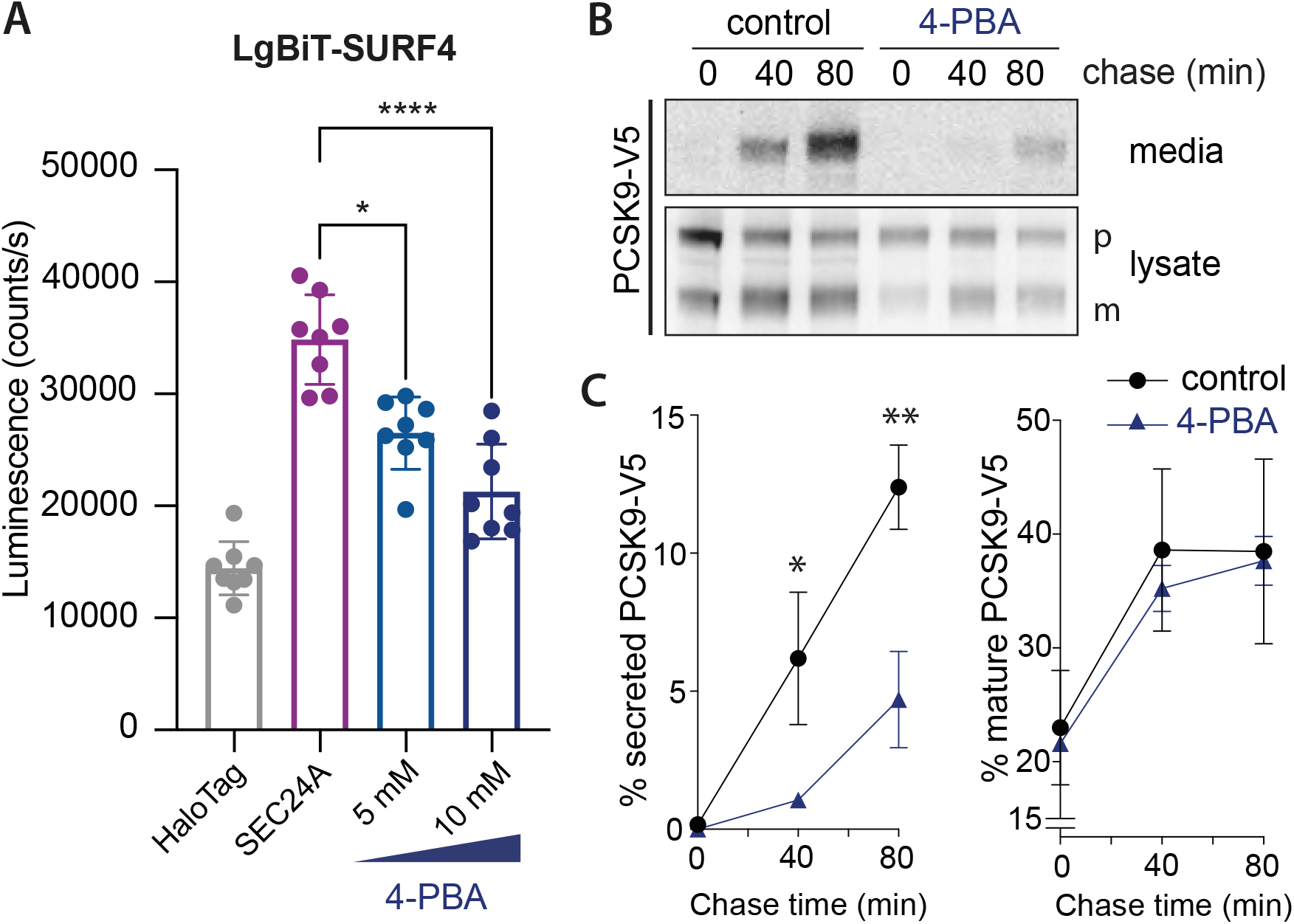
4-PBA reduces the interaction between SEC24A and SURF4 and impairs PCSK9 secretion. **(A)** Luminescence that reports on the interaction between SURF4 and SEC24A was measured in the presence of the indicated concentrations of 4-PBA. The NanoBiT reporters were induced overnight and cells were treated with 4-PBA for 4h. The graph shows the mean luminescence ± SD (n = 8); statistical test was a one-way ANOVA with Dunnett’s correction for multiple comparisons, * p<0.05, ** p<0.01, *** p<0.001, **** p<0.0001. **(B)** PCSK9-V5 maturation and secretion was examined in the presence of 10mM 4-PBA by pulse-chase with [^35^S]methionine. PCSK9-V5 was immunoprecipitated with α-V5 from lysates and the conditioned media at the indicated times and detected by SDS-PAGE and autoradiography. **(C)** PCSK9 secretion into the media (left plot) and maturation (right plot) were quantified by phosphorimage analysis as described for Figure 1. Plots show the mean ± SD of three independent experiments. Statistical test was an unpaired t-test, * p<0.05, ** p<0.01.

### *In vivo* proteomic analysis of *SURF4 KO* cells reveals new clients

We next aimed to more broadly survey how secretion might be impacted by perturbation of the SEC24A-SURF4 interaction. We therefore sought to identify additional secretory proteins that require SURF4 to exit the ER. We used a semi-quantitative proteomic approach to profile the secreted proteins in wild-type and SURF4-deficient (*SURF4KO*) HEK TREx-293 and HuH7 cells. Wild-type and *SURF4KO* cells were labelled with either heavy or intermediate isotopes of arginine and lysine, including label-switch biological duplicates. After the third passage, cells were starved of methionine and then labelled with L-azidohomoalanine (AHA), a methionine analog that enables purification of nascent proteins using click-chemistry. Conditioned media was collected, AHA-containing proteins purified by alkyne-agarose affinity chromatography, and proteins identified by mass spectrometry. A total of 344 and 171 proteins were identified in HEK-293 and HuH7 media fractions, respectively, with good correlation between replicate experiments (Figure 4A, B; Figure S4A, B). Deletion of SURF4 resulted in significantly reduced secretion of 10 proteins in HEK-293 cells and 18 in HuH7 cells (Figure 4C, D, Table S1). Consistent with previous observations, NUCB1 and NUCB2 were among the top hits affected in *SURF4KO* cells (Huang *et al*, 2021). Nucleobindins, together with Cab45, overlap as SURF4-dependent hits between the two cell lineages (Figure 4E). Many SURF4-dependent cargo proteins contain ER-ESCAPE motifs immediately downstream of the predicted signal peptide cleavage sites (Figure S4C). SERPINE1, a protein whose secretion was affected in HuH7 cells, and two HEK-293 hits (FBN2 and LTBP1) do not possess ER-ESCAPE motifs but instead contain a Cardin-Weintraub (CW) motif which has also been shown to mediate cargo-SURF4 interaction (Tang et al., PNAS in press). Other prominent shared features of SURF4 clients include Ca-binding and propensity for oligomerization (Figure S4C).

**Figure 4.**
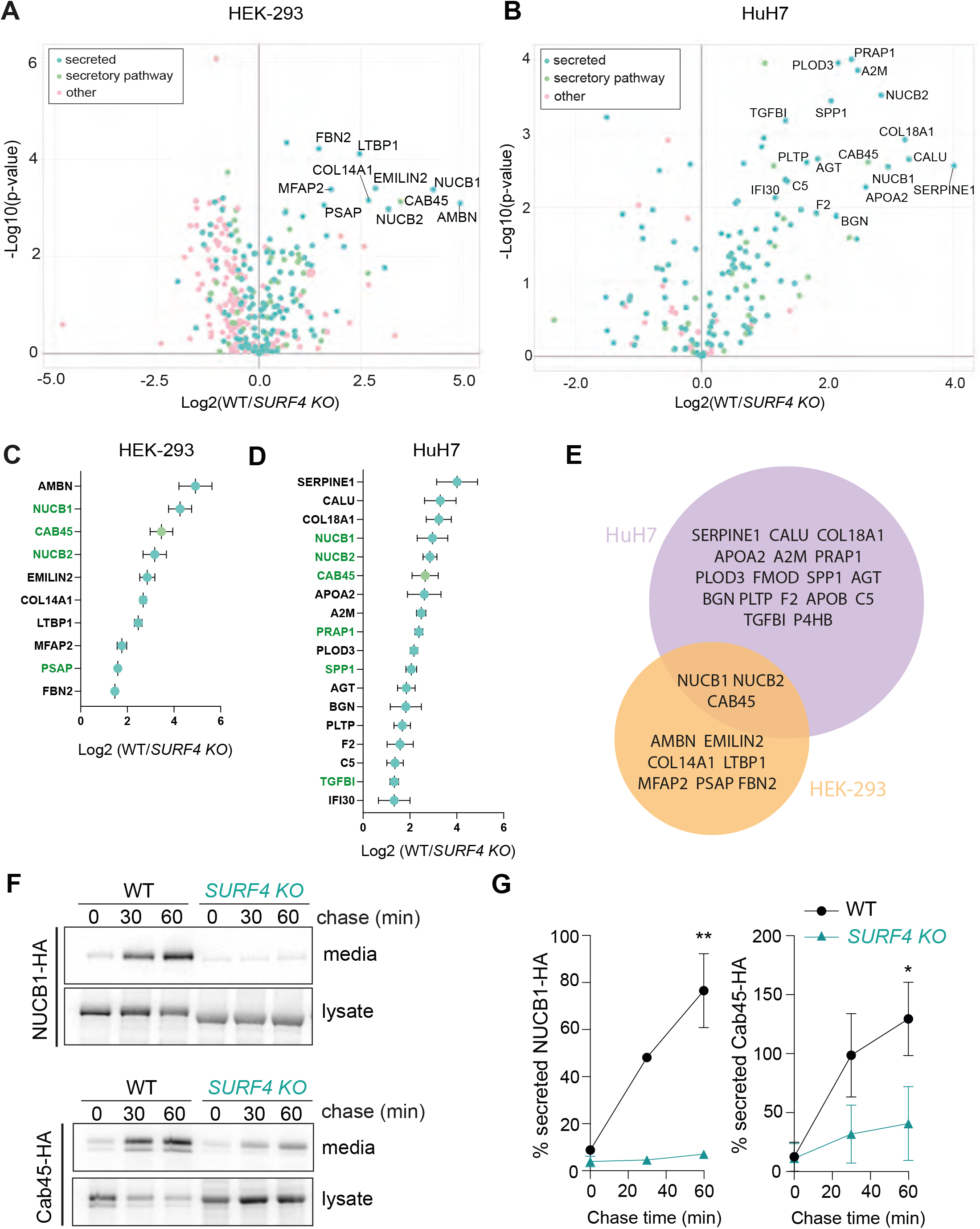
Proteomic quantification of secretion defects in SURF4 KO cells. **(A)**,(B)Volcano plot of SILAC-based quantification of proteins identified in the conditioned media of HEK-293 TREx or HuH7 WT or *SURF4 KO* cells. The proteins indicated with names have FDR-adjusted *p*<0.05 and log2 fold-change >1.25 **(C), (D)** Proteins significantly depleted in the HEK-293 TREx or HuH7 *SURF4* KO cell lines were ranked according to their degree of depletion. Proteins coloured green contain ER-ESCAPE motifs at their predicted N-termini. **(E)** Venn diagram of HEK-293 TREx and HuH7 top hits. **(F)** NUCB1-HA and Cab45-HA secretion was quantified in WT and *SURF4* KO cell lines by pulse chase with [^35^S]methionine. NUCB1 or Cab45 were immunoprecipitated with α-HA from lysates and conditioned media at the indicated times and detected by SDS-PAGE and autoradiography. Autoradiographs are representative of three independent experiments. **(G)** Quantification of NUCB1 or Cab45 secretion by phosphorimage analysis: plots show the mean ± SD of three independent experiments. Statistical test was unpaired t-test, * p<0.05, ** p<0.01.

We validated two SURF4 clients identified in our proteomic analysis using pulse-chase experiments. We measured secretion of HA-tagged NUCB1 and Cab45 in wild-type and *SURF4KO* cells. Indeed, *SURF4KO* cells showed a significant reduction in release of NUCB1-HA and Cab45-HA to the media, consistent with defects in secretion (Figure 4C, D). Secretion of NUCB1-HA could be partially rescued by transient transfection with plasmid-borne SURF4 cDNA (Figure S5).

### NUCB1, Cab45 and PCSK9 use an ER-ESCAPE motif for ER export

To confirm the dependence of SURF4 clients on their ER-ESCAPE motifs (Figure 5A), we mutated the ϕ-P-ϕ residues to glutamic acid, which had previously been shown to impair ER export (Yin *et al*, 2018). Replacement of the ER-ESCAPE motifs in NUCB1 and Cab45 with an EEE peptide caused significant secretion defects (Figure 5B, C). We note that Cab45 with the mutated ER-ESCAPE motif migrated at a lower molecular weight than the wild-type protein (Figure 5B), suggesting impaired glycosylation associated with this mutation. To rule out indirect effects of glycosylation defects perturbing secretion, we mutated the N-glycosylation acceptor site in Cab45 (Scherer *et al*, 1996), Asn40 to Gly (Cab45-N40G). This mutant migrated at a similar apparent molecular weight as Cab45-EEE but was secreted from cells relatively efficiently (Figure S6), confirming that glycosylation defects *per se* cannot explain the intracellular retention of Cab45-EEE.

**Figure 5:**
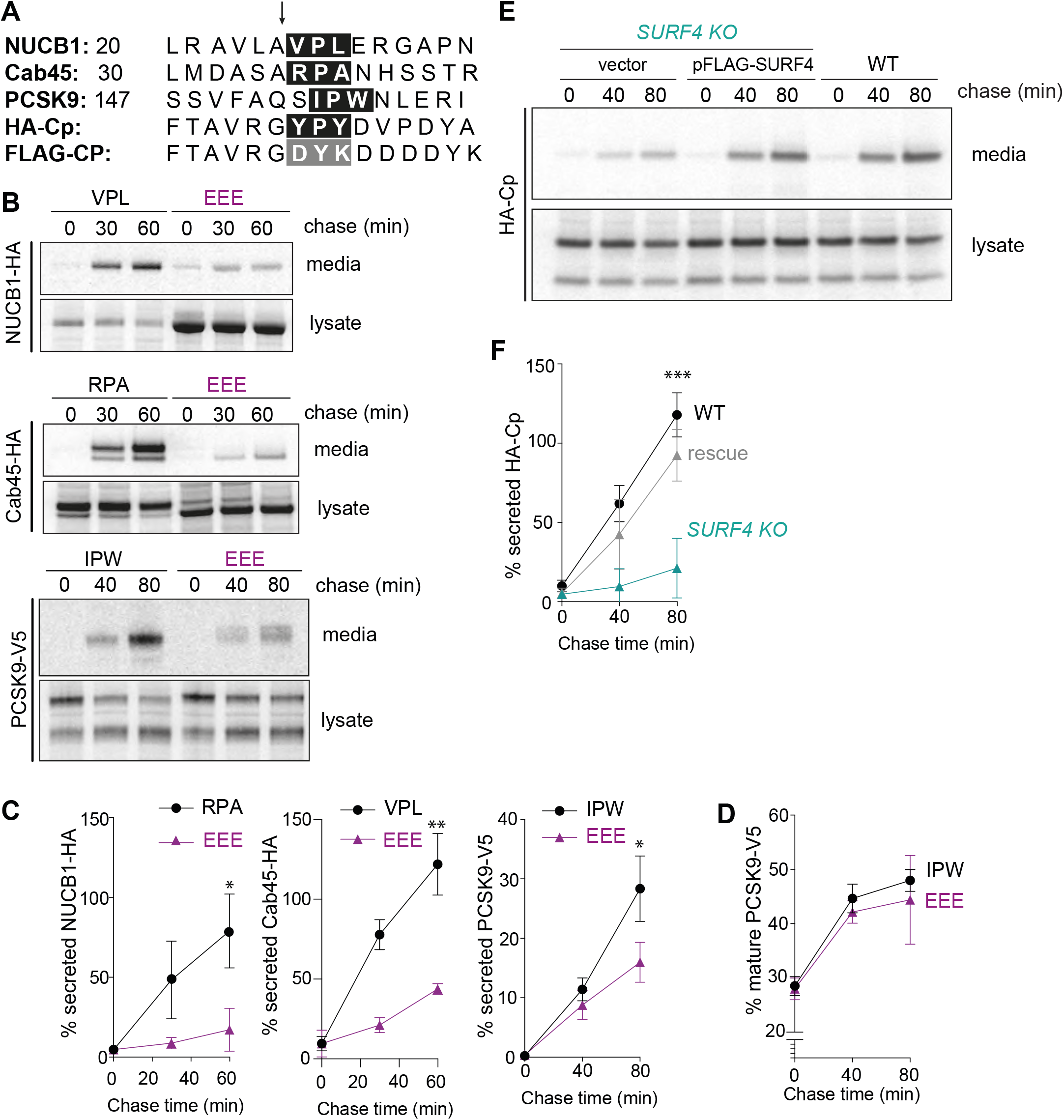
ER-ESCAPE motifs are necessary and sufficient for SURF4-mediated export. **(A)** N-terminal sequences of 3 *bona fide* SURF4 clients (NUCB1, Cab45 and PCSK9) along with a heterologous SURF4 cargo client (HA-Cp), and a bulk flow marker (FLAG-Cp) that lacks an ER-ESCAPE motif. The arrow indicates the site of signal peptide or propeptide cleavage, and the ER-ESCAPE motif is highlighted in the black box. **(B)** Secretion of NUCB1-HA, Cab45-HA and PCSK9-V5 and their corresponding ER-ESCAPE mutants was examined by pulse-chase. **(C)** Percentage secretion was quantified by phosphorimage analysis; plots show the mean ± SD of three independent experiments. Statistical test was unpaired t-test, * p<0.05, ** p<0.01, *** p<0.001. **(D)** PCSK maturation was quantified by phosphorimage anaylsis as described in Figure 1B. **(E)** HA-Cp secretion was examined by pulse-chase in WT, *SURF4 KO*, and *SURF4 KO* cells complemented with a FLAG-SURF4 rescue plasmid. HA-Cp was immunoprecipitated with α-HA from lysates and conditioned media at the indicated times and detected by SDS-PAGE and autoradiography. **(F)** Percentage of secreted HA-Cp was quantified by phosphorimage analysis. Plots show the mean ± SD of three independent experiments. Statistical test was a one-way ANOVA with Dunnett’s correction for multiple comparisons, *** p<0.001.

PCSK9 undergoes auto-catalytic cleavage within the ER such that a propeptide region is cleaved, revealing a new N-terminus. This processing is required for ER export, and yields a ϕ-P-ϕ motif at the +1 position following cleavage (Figure 5A). We tested whether glutamic acid substitutions in this putative ER-ESCAPE motif impaired PCSK9 secretion. Indeed, PCSK9-EEE was markedly reduced in its secretion efficiency, whereas proteolytic processing remained intact (Figure 5B-D). Together, these data support the model that SURF4 clients, including PCKS9, use ER-ESCAPE motifs that are revealed either by signal peptide or propeptide cleavage.

Having demonstrated that the ER-ESCAPE motif is necessary for secretion of several SURF4 clients, we next tested whether this signal is sufficient to direct SURF4-mediated secretion. We turned again to the bulk flow marker, Cp, using an HA-tagged form that exposes a YPY tripeptide after signal peptide cleavage (Figure 5A). Unlike FLAG-Cp (Figure 1C), which reveals DYK at the N-terminus (Figure 5A), HA-Cp secretion was dependent on SURF4, being reduced in the *SURF4KO* line, and rescued by reintroduction of SURF4 cDNA (Figure 5E, F). These findings support the model that N-terminal ϕ-Pro-ϕ signals are necessary and sufficient to drive secretion via interaction with SURF4.

### Selective chemical inhibition of SURF4 client secretion

We next tested whether SURF4 clients were differentially impaired in their secretion by 4-PBA-mediated occlusion of the SEC24 B-site. Similar to the effects observed for PCSK9, secretion of NUCB1-HA and Cab45-HA was reduced in the presence of 4-PBA (Figure 6A). In addition, secretion of the heterologous protein HA-Cp, which carries an ER-ESCAPE motif, was also reduced by the presence of the small molecule, albeit to a lesser extent than PCKS9, NUCB1 or Cab45 (Figure 6C). In contrast, the bona fide bulk flow marker, FLAG-Cp, that lacks the SURF4 export signal was unaffected (Figure 6C, D).

**Figure 6:**
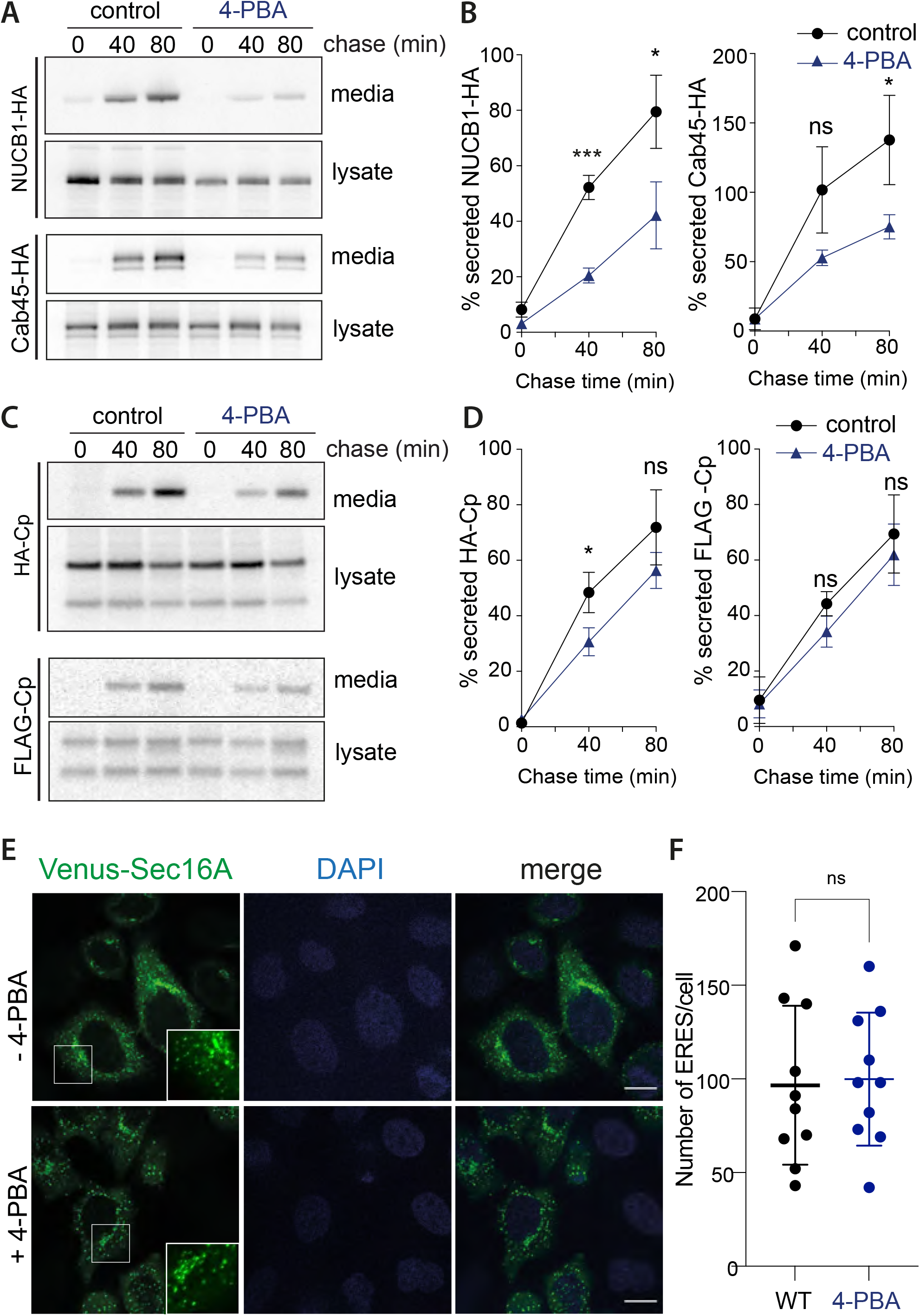
4-PBA selectively inhibits secretion of SURF4 clients but not a *bona fide* bulk flow marker. **(A, C)** Secretion of the indicated proteins was examined in the presence of 10mM 4-PBA by pulse-chase with [^35^S]methionine. NUCB1-HA, Cab45-HA, and HA-Cp were immunoprecipitated with α-HA from lysates and conditioned media at the indicated times and detected by SDS-PAGE and autoradiography. For FLAG-Cp α-FLAG was used for immunoprecipitation **(B, D)** Percentage secretion into the media was quantified by phosphorimage analysis. Plots show the mean ± SD of three independent experiments. Statistical test was an unpaired t-test, * p<0.05, ** p<0.01, *** p<0.001. **(E)** HeLa cells stably expressing Venus-Sec16A were fixed, stained with DAPI and subjected to confocal microscopy. Scale bars, 10 μm. **(F)** Quantification of the number of ERES in the indicated strains expressing Venus-Sec16. Values represent the number of Sec16-positive puncta identified in a single confocal image plane in the center of a cell. A total of 10 cells from each condition was measured. Error bars represent SD; statistical test was an unpaired t-test.

Despite the lack of effect on bulk secretion, we were concerned that 4-PBA might perturb secretion more broadly. Protein export from the ER generally occurs via discrete ER-exit sites (ERES) that are sites of cargo concentration and vesicle formation. To further explore the possibility of a broader effect of 4-PBA on protein export we examined the localisation of the COPII accessory component, Sec16A, which marks ERES in cells (Watson *et al*, 2006), in the presence of 4-PBA. Using confocal microscopy, we observed that when HeLa cells expressing Venus-tagged Sec16A were treated with 10 mM 4-PBA, the structure and number of ERES was unaffected (Figure 6E, F). These results suggest that the action of 4-PBA is specifically linked to its occlusion of the SEC24 B-site. In order to investigate the broader effects of 4-PBA in secretion, we performed secretome analysis using SILAC and AHA labelling. After passage in media containing either light or heavy arginine and lysine isotopes, WT HEK-293 cells were labelled with AHA or AHA plus 10 mM 4-PBA for 4h. Media was collected, and secreted proteins recovered and quantified as described above. Secretion of relatively few proteins was reduced upon treatment with 4-PBA, with SURF4 clients the most prominent proteins affected (Figure S7A, B). Most secretory proteins detected were not affected by 4-PBA treatment (Figure S7C).

## Discussion

Selective ER export relies on a series of protein-protein interactions that specify loading of secretory proteins into nascent transport vesicles. At the top of this interaction pyramid is Sec24, the adaptor protein of the COPII coat (Geva & Schuldiner, 2014). All organisms have multiple Sec24 isoforms, and each isoform likely uses multiple sites of interaction to drive binding of diverse secretory cargo and cargo receptors. This hierarchical arrangement ensures capture of the full spectrum of protein sequence and structure that uses this pathway (Adolf *et al*, 2019; 2016; Bonnon *et al*, 2010; Mancias & Goldberg, 2008) (Mancias & Goldberg, 2007). In yeast, four different cargo-binding sites, referred to as A, B, C, and D-sites, on Sec24 allow recognition of a diverse set of sorting signals (Miller *et al*, 2003; Mossessova *et al*, 2003; Pagant *et al*, 2015). Proteomics of vesicles generated with different mammalian Sec24 isoforms revealed both shared and isoform-specific clients (Adolf *et al*, 2019). The nature of specialized cargo-binding events for distinct Sec24 binding sites raises the appealing possibility of selective inhibition of secretion, an otherwise essential process. Here, we have mapped the interaction network responsible for secretion of the cholesterol regulator, PCSK9. We confirm that SEC24A and the cargo receptor, SURF4, drive PCSK9 secretion, and demonstrate a role for the B-site of SEC24A in this process. A known small molecule, 4-PBA, which binds the SEC24 B-site (Ma *et al*, 2017) inhibits the SEC24A-SURF4 interaction and impairs secretion of PCSK9 and other SURF4 clients. This establishes the principle that small molecule modulation of cargo selection can selectively alter the cellular secretome.

4-PBA was first described as a chemical chaperone as thought to improve ER folding capacity (Welch & Brown, 1996). Generally, 4-PBA suppresses cellular responses against the accumulation of misfolded proteins in the ER, such as the unfolded protein response (UPR), in both yeast and mammalian cells (Kubota *et al*, 2006; Ozcan *et al*, 2006; Pineau *et al*, 2009; Mai *et al*, 2018), however the mechanism of action of 4-PBA was unknown. Recently it has been described that 4-PBA occludes cargo recognition sites in the COPII coat generating cargo-packaging defects that result in increased capture of ER resident proteins (Ma *et al*, 2017). That observation explains the previously described effects of 4-PBA facilitating trafficking of misfolded proteins (Balch *et al*, 2008; Liu *et al*, 2004). 4-PBA mimics the structure of C-terminal hydrophobic sorting signals that interact with the SEC24 B-site (Ma *et al*, 2017). This relatively broad-spectrum effect of 4-PBA is consistent with the conserved nature of the Sec24 B-site across all SEC24 isoforms, and highlights its limited utility as a selective therapeutic. However, structural studies have shown that subtle changes in the vicinity of the B-site of human SEC24C and SEC24D results in a marked specificity for distinct export signals (Mancias & Goldberg, 2008), raising the prospect that more refined small molecules might be able to selectively target different B-site isoforms. Indeed, understanding the molecular basis for SEC24A discrimination of SURF4 is an important question that should inform additional chemical approaches.

In addition to the SEC24-SURF4 interaction, secretion of PCSK9 and other SURF4-dependent cargoes relies on interaction between an unknown SURF4 lumenal domain and the ER-ESCAPE motifs of its clients (Yin *et al*, 2018). This interaction represents another potentially druggable interface that might be similarly targeted by small molecule inhibition. Since multiple diverse cargo proteins interact with the Sec24 B-site (Miller *et al*, 2003; Mossessova *et al*, 2003; Mancias & Goldberg, 2008; Ma *et al*, 2017), perturbing SURF4-client interaction may represent a more tractable way forward to therapeutic intervention. However, the client repertoire of SURF4 seems substantial (Yin *et al*, 2018; Ordonez *et al*, 2021; Huang *et al*, 2021), and being able to selectively inhibit a subset of receptor-cargo interactions may be difficult. Nonetheless, like SEC24, SURF4 also appears to be a multivalent export facilitator, since it also recognises a second ER export signal, the Cardin-Weintraub (CW) motif, found on Shh (Tang et al., in press). Discerning which SURF4 clients use the ER-ESCAPE versus CW motifs, and defining the sites on SURF4 that mediate these interactions is another important open question that should inform future small molecule screening approaches. We also note that multiple SURF4 clients are Ca-binding proteins and/or undergo oligomerization, consistent with previous observations that suggested that SURF4 is important to (i) prevent premature oligomerization within the ER; and (ii) rapidly remove these proteins from the Ca-rich ER lumen before inappropriate oligomerization (Yin *et al*, 2018). Whether a putative chaperone-like function is also mediated by the ER-ESCAPE (or CW) motif, or is instead conferred by a separate domain of SURF4 remains to be determined. Although no obvious induction of the UPR has been observed upon the loss of Surf4 (Huang *et al*, 2021; Wang *et al*, 2021; Ordonez *et al*, 2021). Dissecting cargo-binding and chaperone functions of SURF4, and understanding how interactions at these sites on SURF4 inform the interaction with SEC24 will be an important priority.

Our findings specifically raise the prospect of selective inhibition of PCSK9 secretion. In recent years, PCSK9 has emerged as a key therapeutic target for lowering circulating LDL levels. PCSK9 monoclonal antibodies not only markedly lower LDL levels but also prevent cardiovascular events. However, PCSK9 monoclonal therapeutics require monthly injections, and their manufacturing process is expensive. Therefore, non-antibody treatments to inhibit PCSK9 function are actively being developed as alternative therapies (Shapiro *et al*, 2018; Nishikido & Ray, 2018). Our work here establishes that small-molecule inhibition of the secretory pathway may be a useful alternative to inhibit PCSK9 function. Indeed, 4-PBA has been shown to reduce atherosclerosis in mice (Huang *et al*, 2017; Lynn *et al*, 2019), although the molecular basis for this effect was not known. Finally, given the wealth of structural information on cargo selection by SEC24, additional small molecule screening based on the types of protein-protein interaction assays described here suggests a path forward for more specific inhibition of protein secretion, and raises the possibility for systemic modulation other health-related secretory proteins.

## Acknowledgements

We thank David Stephens for the Venus-Sec16 HeLa cell line, Julia von Blume for Cab45 reagents, Ari Helenius for the HA-Cp bulk flow reporter, and members of the Miller lab for constructive feedback on the project. This project is supported through a research collaboration between AstraZeneca UK Limited and the Medical Research Council, reference BSF29 (to EAM and DG), by the Medical Research Council under award number MRC_UP_1201/10 (to EAM) and by the National Institutes of Health (R01GM117473 to MB).

## Materials and Methods

### Cell lines

HEK TREx-293 and HuH7 cell lines used in this study were cultured in Dulbecco’s Modified Eagle’s Medium (DMEM, Gibco) with 10% fetal bovine serum (Qualified FBS, Gibco). HEK TREx-293 SURF4 KO was previously described (Huang *et al*, 2021). For CRISPR-Cas9 mediated knockout of Sec24A and Sec24B, guide RNAs targeting exon 1 (5’-GGCCCAGAACGGAGCCGCCT-3’) of SEC24A and exon 3 of SEC24B (5’-CGTATCCTAGTGTTTCATAT-3’) were designed using the CRISPR design tool at (https://benchling.com) and cloned into pX458 (pSpCas9 BB-2A-GFP, Addgene plasmid # 48138). Multiple single-cell derived knockout clones per each guide RNA were isolated and screened for the gene disruption by western blotting. SEC24C and SEC24D KO cell lines were described previously (Bisnett *et al*, 2021). Cells were validated by immunoblotting or quantitative RT-PCR (Figure S8). ERES were visualized using a HeLa cell line expressing Venus-tagged Sec16A (Watson *et al*, 2006), a kind gift from Dr David Stephens. Transient transfections were performed with TransIt 293 reagent from *Mirus*. All cell lines were routinely checked for mycoplasma contamination and tested negative.

### Constructs and reagents

The plasmids used in this study are listed in Table S2. PCSK9 for mammalian expression in the pcDNA5 vector containing a C-terminal V5 tag was generated by GeneScript. PCSK9-EEE mutant contained three-point mutations within the ER-ESCAPE motif: I154E, P155E, W156E and was generated using the *QuikChange Lightning Multi Site-Directed Mutagenesis Kit (Agilent Technologies)*. NUCB1 for mammalian expression in the pcDNA3.1 vector containing a C-terminal HA tag was generated by GeneScript. The NUCB1-EEE mutant contained three-point mutations within the ER-ESCAPE motif: V27E, P28E, L29E and was generated using the QuikChange Lightning Multi Site-Directed Mutagenesis Kit (Agilent Technologies). The Cab45 plasmid for mammalian expression in the pLPCX vector containing a C-terminal HA tag was kindly provided by Julia Von Blume (Von Blume *et al*, 2012). The Cab45-EEE mutant contained three-point mutations within the ER-ESCAPE motif: R37E, P38E, A39E and was generated using the *QuikChange Lightning Multi Site-Directed Mutagenesis Kit (Agilent Technologies)*. SURF4 for mammalian expression in the pcDNA3.1 vector containing a N-terminal FLAG tag was generated by GeneScript. The HA-Cp construct in the pSV-SPORT1 (Thor *et al*, 2009) was kindly provided by Ari Helenius. The FLAG-tagged version of Cp was created by sequential mutagenesis reactions using the QuikChange Lightning Multi Site-Directed Mutagenesis Kit (Agilent Technologies).

HaloTag, SEC24A, SEC24C and SURF4 plasmids for the NanoBiT PPI assays were in the pcDNA3.1 vector containing the SmBiT tag for HaloTag, SEC24A and SEC24C and the LgBiT tag for SURF4. The SEC24A ORF was amplified by PCR from a commercially available plasmid (cDNA SEC24A in pcDNA3.1+/-C-(K)-DYK, GeneScript) and subcloned into the pFC36K-SmBiT and pFN35k-SmBiT backbones including the TK promoter (*Promega*). SURF4 ORF was amplified by PCR from a commercially available plasmid (cDNA SURF4 in pcDNA3.1+/-C-(K)-DYK, GeneScript) and subcloned into the pFC34-LgBiT and pFN33-LgBiT backbones including the TK promoter (*Promega*). These constructs resulted in very low expression of SEC24A and SURF4 under the TK promoter. The fragments containing SmBiT-SEC24A or SEC24A-SmBiT and LgBiT-SURF4 or SURF4-LgBiT were PCR amplified and subcloned into the HindIII/NotI sites of pcDNA3.1. The HaloTag-SmBiT fragment was PCR amplified from the commercially available plasmid *(Promega, Cat no: N202A*) and also subcloned into pcDNA3.1. The SEC24A B- and C-site mutants were generated on pcDNA3.1-SmBiT-SEC24A using QuickChange site-directed mutagenesis (Agilent Technologies).

4-PBA (4-phenylbutyric acid, *Sigma*) and 5-PVA (5-phenylvaleric acid, *Sigma*) were dissolved in equimolar amount of NaOH to prepare a 0.5M stock solution with pH 7.

### Pulse-chase

Cells were starved in Methionine/Cysteine-free DMEM (Gibco, #21013024) for 30 min, pulsed with 15 uCi/mL ^35^Smethionine/cysteine *(EasyTag EXPRESS* ^*35*^*S, Perkin Elmer)* and chased in complete DMEM containing 10% FBS. At each time point media and cells were collected and cells were lysed in Buffer A (50 mM Tris pH 7, 150 mM NaCl, 1% (v/v) Triton X-100, 2 mM EDTA) supplemented with Protease inhibitors (Complete EDTA-free, Roche). Media and cell lysates were pre-cleared and the protein of interest was immunoprecipitated. Radiolabelled immunoprecipitated proteins were eluted in Loading buffer (50 mM Tris pH 6.8, 0.1% (v/v) glycerol, 20% (p/v) SDS, 5% (v/v) b-mercaptoethanol, 1 mg/mL Bromophenol Blue), separated on NuPAGE 4-12% Bis-Tris gels *(ThermoFisher Scientific)* and detected by phosphorimaging using a Typhoon scanner *(GE Healthcare)*. The protein bands were quantified using Fiji and the percentage of the mature or secreted band in each sample was plotted with Prism 7.0 (*GraphPad Software*).

### SILAC-AHA labelling and enrichment of endogenously secreted proteins

In order to elucidate SURF4-dependent secretome, the method described in (Eichelbaum *et al*, 2019) was adapted. To ensure SILAC label incorporation, WT and *SURF4 KO* TREx-293 cells were grown for 3 passages in DMEM (AthenaES, #0420) supplemented with 10% FBS and with either heavy (84 μg/mL L-arginine-HCl, 13C6, 15N4, *Thermo Fischer*, #89990; 145 μg/mL L-lysine 2HCl, 13C6, 15N2, Thermo Fischer, #88209) or intermediate (84 μg/mL L-arginine-HCl 13C6, Thermo Fischer, #88210; 146 μg/mL L-lysine-2HCl, 4,4,5,5,D4, Thermo Fischer, #88437) SILAC labels, supplied with excess L-proline (200 μg/mL, *Formedium* LTD, #DOC0177) and L-methionine (201 μM, AthenaES #0419). Cells at 70% confluency in 10-cm dishes, were starved in SILAC DMEM with no methionine for 30 min and then labelled with L-azidohomoalanine (0.5 mM, AnaSpec, #AS-63669) for 20h. Oppositely labelled media was then combined and treated with 0.5M aminoguanidine-HCl (Sigma #396494), protease inhibitors, and centrifuged to remove remaining cells. Samples, concentrated to ∼250 μL (Merck, Amicon, 3 kDa cut-off), were combined with 250 uL lysis buffer (1% CHAPS, 50 mM HEPES pH 7, 150 mM NaCl, 8M urea) with PIs. 100 uL of alkyne agarose slurry (Jena Bioscience #CLK-1032-2) was washed with IP buffer (1% CHAPS, 50 mM HEPES pH 7, 150 mM NaCl). The washed resin, 2 mM CuSO4 (Chem Cruz #sc-203009A), 2 mM THPTA (Jena Bioscience #CLK-1010-25), and 4 mM sodium ascorbate (Sigma #PHR1279) were added to the samples and these were rotated for 16-20 h at room temperature. The resin was then washed with HPLC-grade water (Fischer #W/016/17), resuspended in SDS buffer (100 mM Tris-HCl pH 8, 1% SDS, 250 mM NaCl, 5 mM EDTA) with 10 mM DTT and incubated at 70° for 15 min. Once cooled, samples were incubated in the dark for 30min with SDS buffer (100 mM Tris–HCl pH 8, 1% SDS, 250 mM NaCl, 5 mM EDTA) with 40 mM iodoacetamide (Sigma, #I1149). The resin was then resuspended in SDS buffer, transferred to a spin column and washed with SDS buffer, 8M urea in 100 mM Tris pH 8, and 20% acetonitrile. The resin was finally resuspended in digestion buffer (100 mM Tris-HCl pH 8, 2 mM CaCl2 and 10% acetonitrile) and processed for mass spectrometry analysis.

### Mass spectrometry and data analysis

Proteins retained in the resin were digested with trypsin (*Promega*) overnight at 37 °C. Post-digestion, samples were centrifuged and each supernatant was transferred to a fresh tube. Then, beads were extracted with 50% acetonitrile/0.5% formic acid (FA) and combined with the corresponding supernatant. Peptide mixtures were phase-reverse fractionated, partially dried in Speed Vac and desalted using home-made C18 (3M *Empore*) stage tip that contained 3 μL porous R3 (*Applied Biosystems)* resin. Bound peptides were eluted sequentially with 30%, 50% and 80% acetonitrile in 0.5% FA and concentrated in a Speed Vac (Savant). The peptide mixtures were desalted and fractionated using self-packed C18 (3M Empore) stage tips filled with 3 μL of porous R3 resin (Applied Biosystems). The stage tips were activated and equilibrated with 50% acetonitrile (MeCN), 80% MeCN/0.5% FA. Peptide mixtures were loaded onto stage tips and washed with 0.5% FA. Then, stage tips were equilibrated with 100 μL of 10 mM ammonium bicarbonate for pH=8 fractionation. Bound peptides were sequentially eluted using 8 × 20 ul of 10 mM ammonium bicarbonate containing increasing concentrations of MeCN and combined into 4 fractions. Eluted peptides were acidified and concentrated in a Savant Speed Vac.

Peptides were then subjected to LC/MS2 analysis. Desalted peptide mixtures were separated using an Ultimate 3000 RSLC nano System *(Thermo Scientific)*, with an acetonitrile gradient, consisting of buffer A (2% MeCN, 0.1% FA) and buffer B (80% MeCN, 0.1% FA) at a flow rate of 300 nl/min for 2h. The HPLC system was coupled to a Q-Exactive Plus hybrid quadrupole-Orbitrap mass spectrometer (ThermoFisher Scientific), equipped with a nanospray ion source. The mass spectrometer was operated in standard data-dependent mode, performed survey full-scan (MS, m/z = 380-1600) with a resolution of 70000, followed by MS2 acquisitions of the 15 most intense ions with a resolution of 17500 and NCE of 27%. MS target values of 1e6 and MS2 target values of 1e5 were used. The isolation window was set as 1.5 m/z and dynamic exclusion for 30s.

The acquired MS/MS raw files were processed using MaxQuant with the integrated Andromeda search engine (v1.6.6.0). MS2 spectra were searched against Homo sapiens Reviewed, UniProt Fasta database (Mar 2019). Carbamidomethylation of cysteines was set as fixed modification, while oxidation of methionine, N-terminal protein acetylation, Met replaced by AHA and reduction of Met replaced by AHA as variable modifications. For AHA-SILAC, Lys0/Arg0 (light), Lys4/Arg6 (intermediate) and Lys8/Arg10 (heavy) were specified as metabolic labels. Enzyme specificity was set to trypsin/p and maximum two missed cleavages were allowed. Protein quantification requires 1 (unique+razor) peptide. Further data visualisation and analysis were carried out using Excel, Perseus (v1.6.2.3) and R (v3.6.1). A protein hit was subjected for further analysis if it had at least 2 SILAC ratios in 4 replicates and >=2 detected peptides. One-sample t-test was used to calculate p-values which were then adjusted for multiple testing with Benjamini-Hochberg method.

### Cell proliferation assays – MTS assays

Cells were seeded into 96-well plates at a density of 8×10^3^ cells/well (100uL) and treated the following day with 5, 10, 20 mM 4-PBA or 5, 10, 20 mM 5-PVA for 4 hours. For the MTS assay, MTS Cell Proliferation Colorimetric Assay Kit (*Generon*) was used following the manufacturer’s instructions. Briefly, 2 hours before the desired time point, 10 μl of the MTS reagent was added into each well and cells were incubated at 37 °C for 2 hours. The absorbance was detected at 490 nm with a Microplate Reader (*Spark plate reader, Tecan*).

### Western Blot analysis

For analysis of total cellular proteins, cells were lysed in 100 mM Tris pH 8.0 with 1% SDS supplemented with protease inhibitor (*Complete EDTA-free, Roche*). Cells in lysis buffer were heated 3 times for 5 min at 95 °C. Protein concentrations were adjusted based on A_280_ values to a final concentration of 2.5 ug/uL and loading buffer was added. 25 ug of proteins were then separated in NuPAGE 4-12% Bis-Tris gels (*ThermoFisher Scientific*), transferred to 0.2 μm nitrocellulose membrane (*Whatman*) and detected with the corresponding antibody. Chemiluminescence was visualized employing HRP-conjugated secondary antibodies and chemiluminescent substrate (*Immobilon Western Chemiluminescent HRP Substrate, Sigma*).

### Confocal microscopy

HeLa venus-Sec16A mRuby-Sec23A, kindly provided by David Stephen’s, were seeded on coverslips in 24-well plates and then treated with or without 10 mM 4-PBA for 4 hours. Then cells were fixed with 4% paraformaldehyde for 30 min and permeabilized with 0.1% Triton X-100 for 10 min and stained with 10 ug/uL DAPI (*Merk*) for 20 min. Coverslips were finally mounted in Prolong Diamond Antifade (*ThermoFisher Scientific*). Images were taken on an Andor Revolution Spinning Disk microscope with a 100x/1.3NA oil immersion objective and an EMCCD camera. Images were collected using the Andor iQ3 software, followed by processing in Fiji to adjust brightness and contrast.

### PPI assays

For PPI luminescence measurements, double KO (*SEC24AKO SURF4KO*) TREx-293 cells were plated at a density of 8 × 10^3^ cells/well (100uL) in DMEM and transfected the following day with the correspondent NanoBiT constructs. 24 hours post-transfection, growth medium was exchanged with Opti-MEM, and cells were incubated for 3.5 hours at 37 °C. *Nano-Glo Live Cell Substrate* (*Promega*) was added, and luminescence was measured using a *Spark plate Reader (Tecan)* at 37 °C.

---

**Figure S1:**
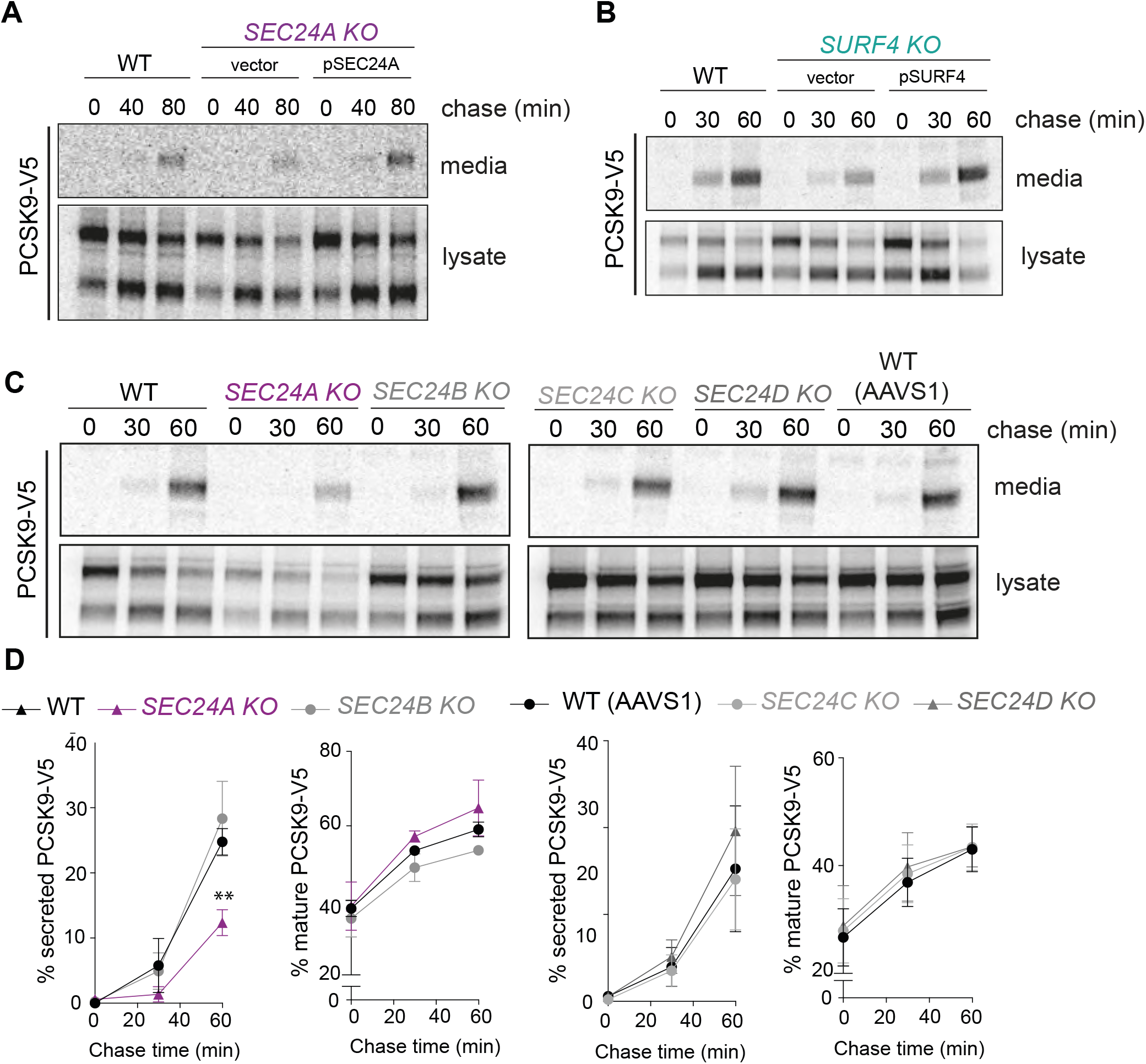
Rescue of PCSK9 secretion defects in SEC24A KO and SURF4 KO cells. **(A)** PCSK9-V5 secretion was examined by pulse-chase in WT, and *KO* cells complemented by transient transfection with the relevant rescue plasmids. PCSK9 was immunoprecipitated with α-V5 from lysates and the conditioned media at the indicated times and detected by SDS-PAGE and autoradiography as described in Figure 1. **(B)** Quantification of PCSK9 secretion shown in A as described in Figure 1. **(C)** PCSK9-V5 secretion was monitored in *SEC24* KO lines indicated, along with the corresponding wild-type lines (HEK TREx293 for *SEC24B* KO and AAVS1 for *SEC24C* and *SEC24D* KO), as described in Figure 1. **(D)** Quantification of PCSK9 secretion shown in C as described in Figure 1.

**Figure S2:**
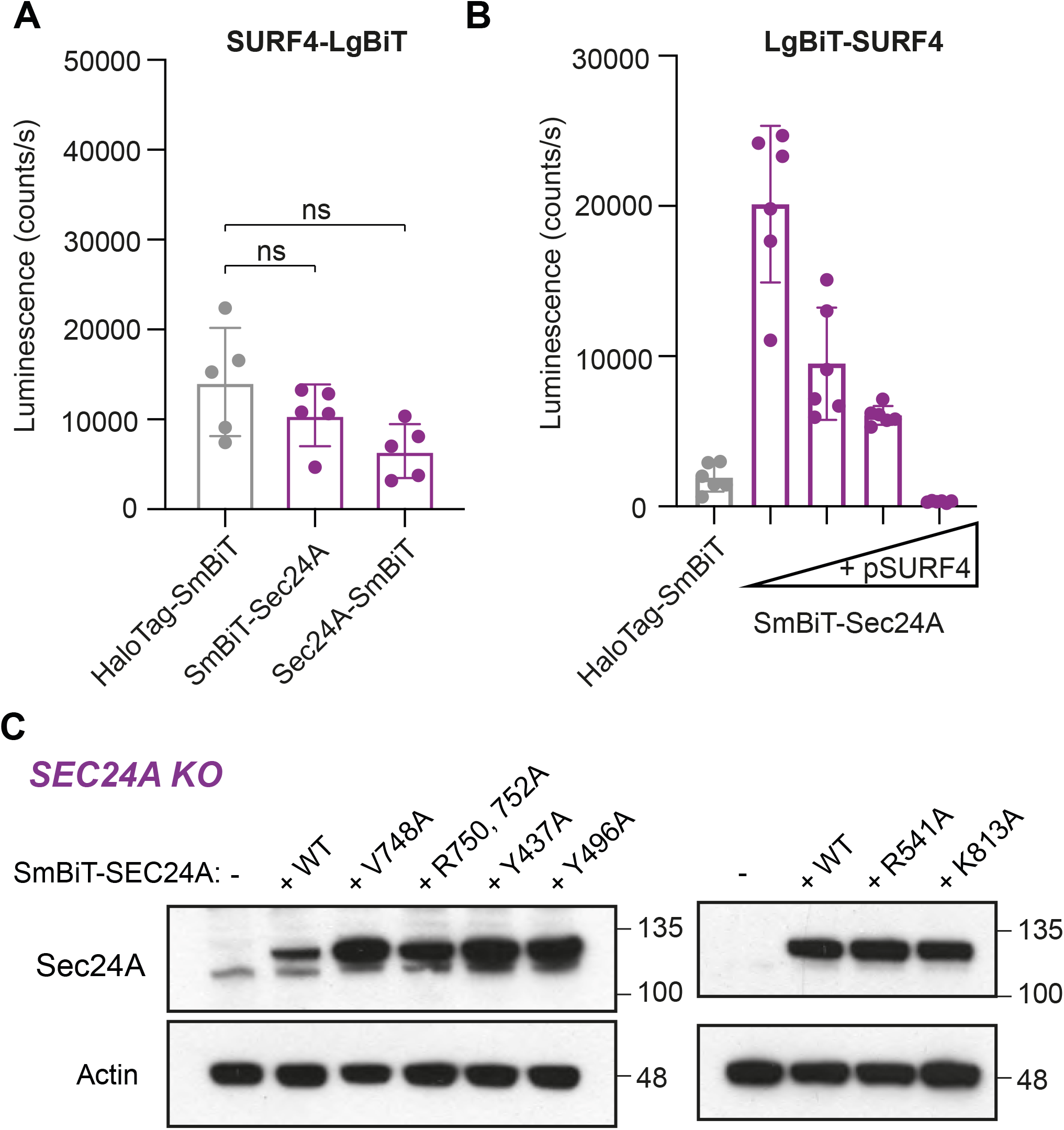
Establishment of the SEC24A-SURF4 NanoBiT assay and stability of SEC24A mutants. **(A)** Control experiments for the NanoBiT assay showing non-optimal tagging orientations and the corresponding negative control in *SEC24A SURF4* double KO cells. The graph shows mean luminescence ± SD (n = 6); statistical test was a one-way ANOVA with Dunnett’s correction for multiple comparisons. **(B)** Specificity of the assay was tested by expressing increasing amounts of untagged fusion partners. Plots show mean ± SD, n=6. **(C)** Stability of B-site (left-hand panel) and C-site (right-hand panel) SEC24A mutants was analysed by immunoblotting cell lysates prepared from equal numbers of cells with α-SEC24A antibodies.

**Figure S3:**
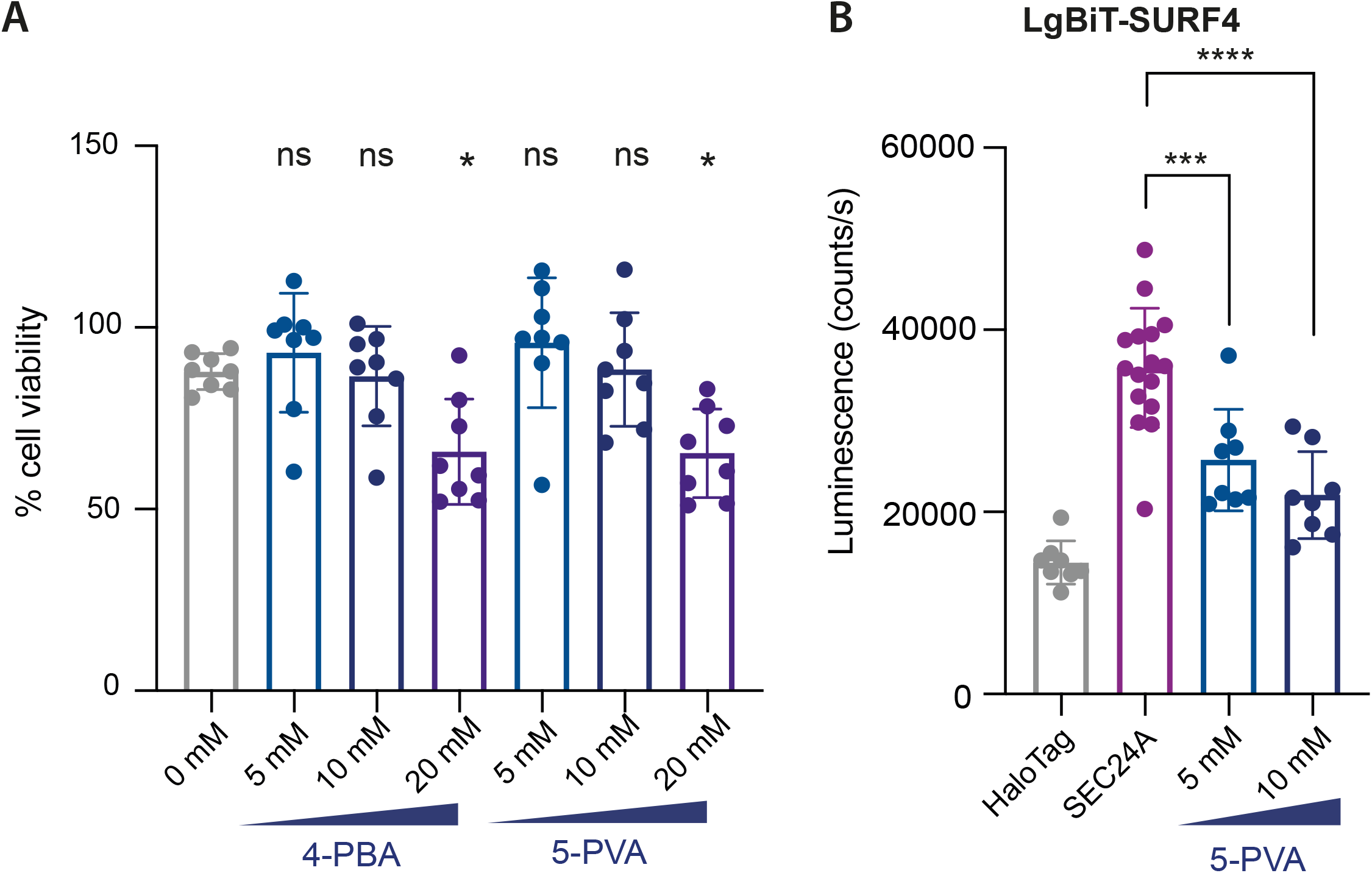
Effects of 4-PBA on cell viability. **(A)** Dose-dependent effects of 4-PBA on cell viability. Cells were grown on plates overnight and then treated for 4h with the indicated concentrations of 4-PBA. Cell viability was measured using the MTS Cell Proliferation Colorimetric Assay Kit. Values are given as mean ± SD, n=8. **(B)** Luminescence that reports on the interaction between SURF4 and SEC24A was measured in the presence of the indicated concentrations of 5-PVA. The NanoBiT reporters were induced overnight and cells were treated with 5-PVA for 4h. The graph shows the mean luminescence ± SD (n = 8; n= 16 for 0 mM 4-PBA); statistical test was a one-way ANOVA with Dunnett’s correction for multiple comparisons, * p<0.05, **** p<0.0001.

**Figure S4:**
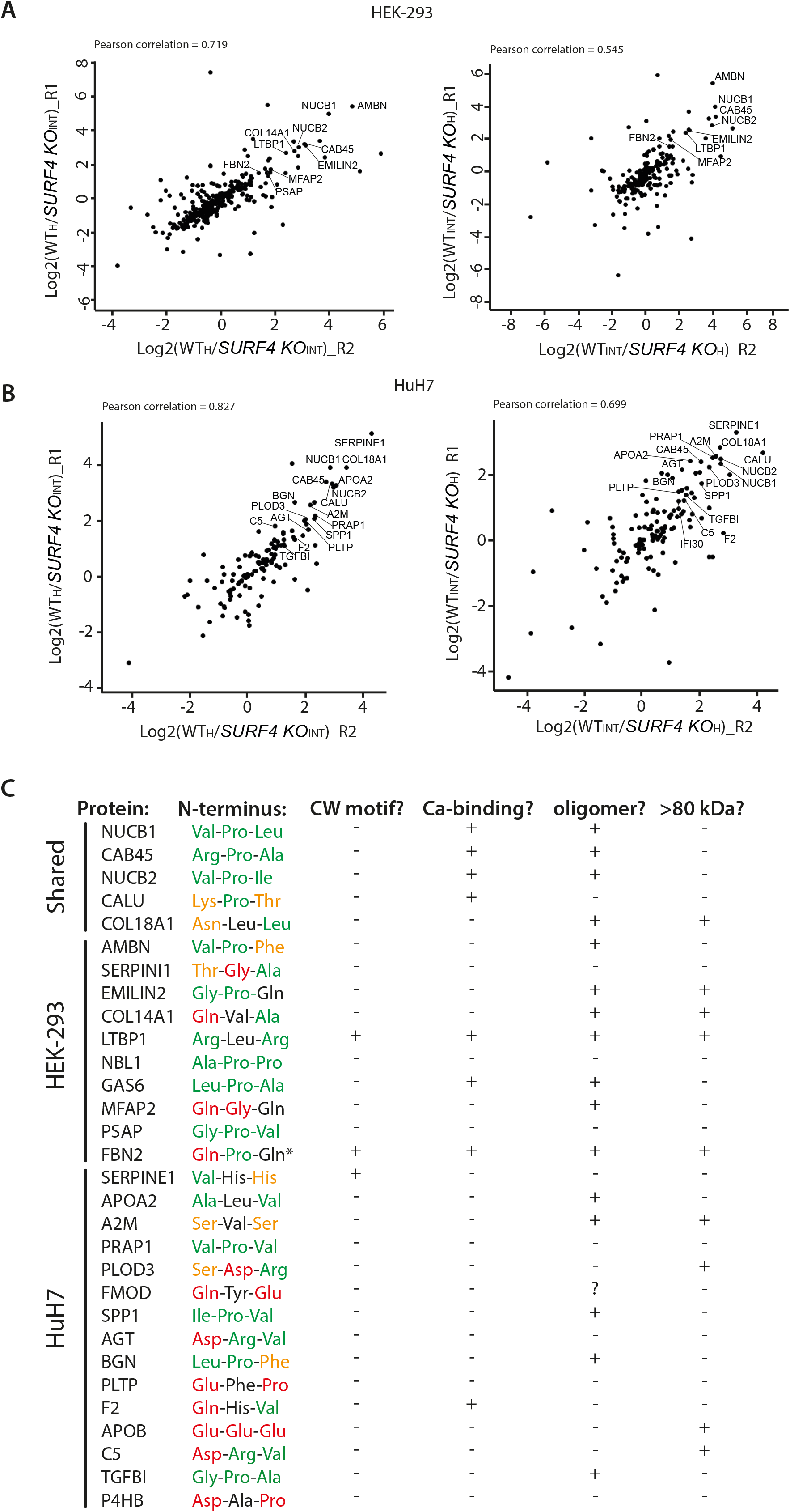
ER-ESCAPE motif sequence of the top SURF4 cargo clients identified in the proteomic quantification. **(A), (B)** Scatter plots of Log2 SILAC ratios of the two replicates within each label-switch pair in HEK-293 or HuH7 cell media samples. Correlation is indicated by Pearson’s correlation coefficient. **(C)** Top SURF4 client hits identified by proteomic analysis and annotated for (i) N-terminal sequence color coded based on relative contribution of each amino acid position to the strength of the ER-ESCAPE motif as described in (Yin et al., 2018) (green = very good, yellow = good, black = neutral, and red = bad); (ii) presence of CW motif; (iii) presence of a Ca-binding domain or annotation as a Ca-binding protein; (iv) propensity to oligomerize; (v) protein size (small/medium <80kDa, large >80 kDa).

**Figure S5:**
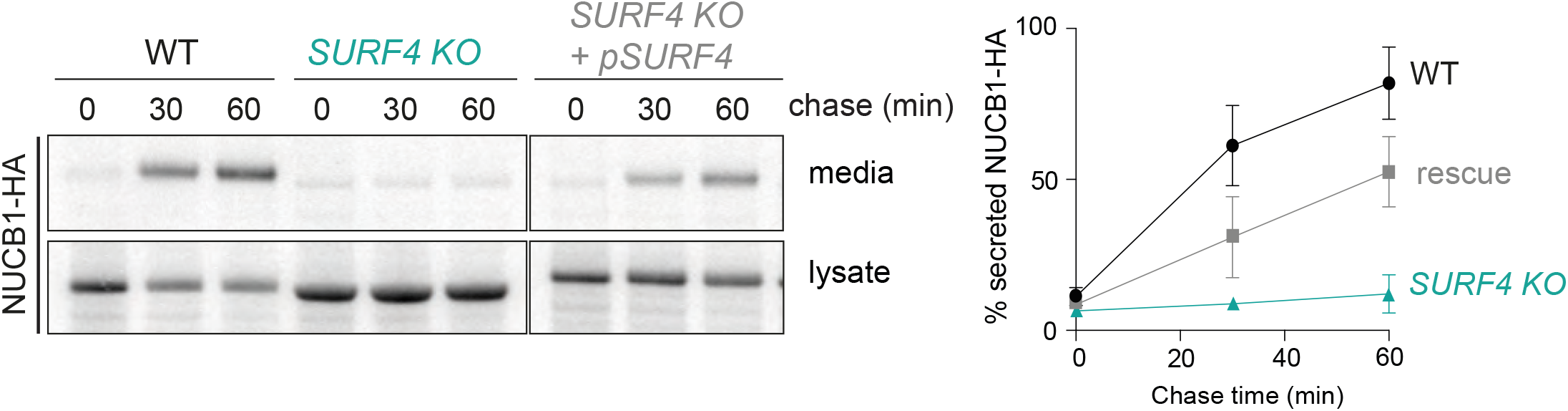
Rescue of NUCB1 secretion defects in SURF4 KO cells. NUCB1-HA secretion was examined by pulse-chase in WT, *SURF4 KO*, and *SURF4 KO* cells complemented with the pFLAG-SURF4 plasmid. NUCB1-HA was immunoprecipitated with α-HA antibodies from lysates and the conditioned media at the indicated times and detected by SDS-PAGE and autoradiography.

**Figure S6:**
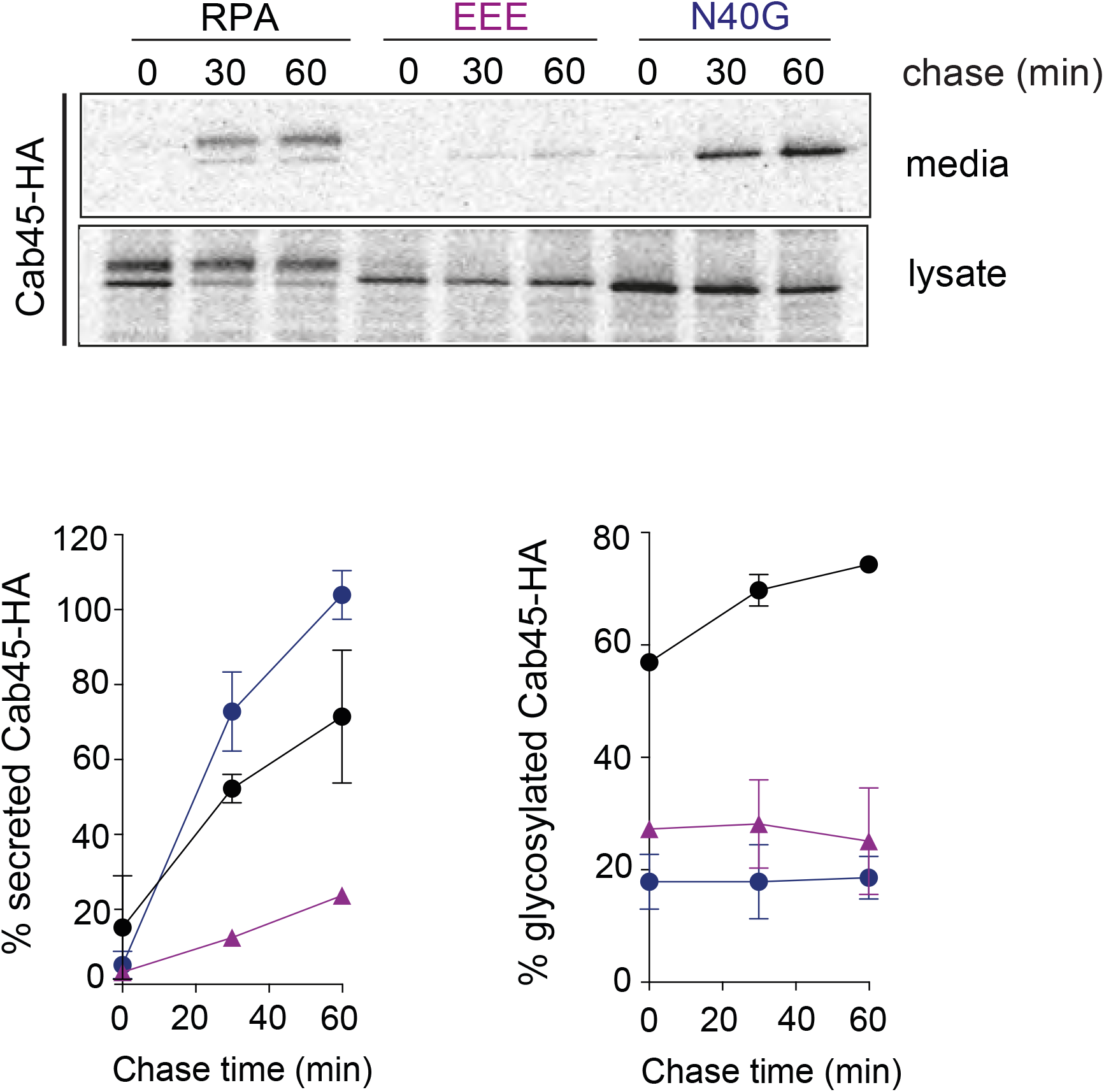
Glycosylation defects do not influence Cab45 secretion. **A)** Cab45-HA secretion was examined by pulse-chase in cells expressing WT (RPA), ER-ESCAPE mutant (EEE) and glycosylation mutant (N40G) variants. Cab45-HA was immunoprecipitated with α-HA antibodies from lysates and the conditioned media at the indicated times and detected by SDS-PAGE and autoradiography. **(B)** Percentage secretion into the media was quantified by phosphorimage analysis. Percentage of glycosylation was quantified as [glycosylated signal / [total protein (glycosylated + unglycosylated). Plots show the mean ± SD of three independent experiments.

**Figure S7:**
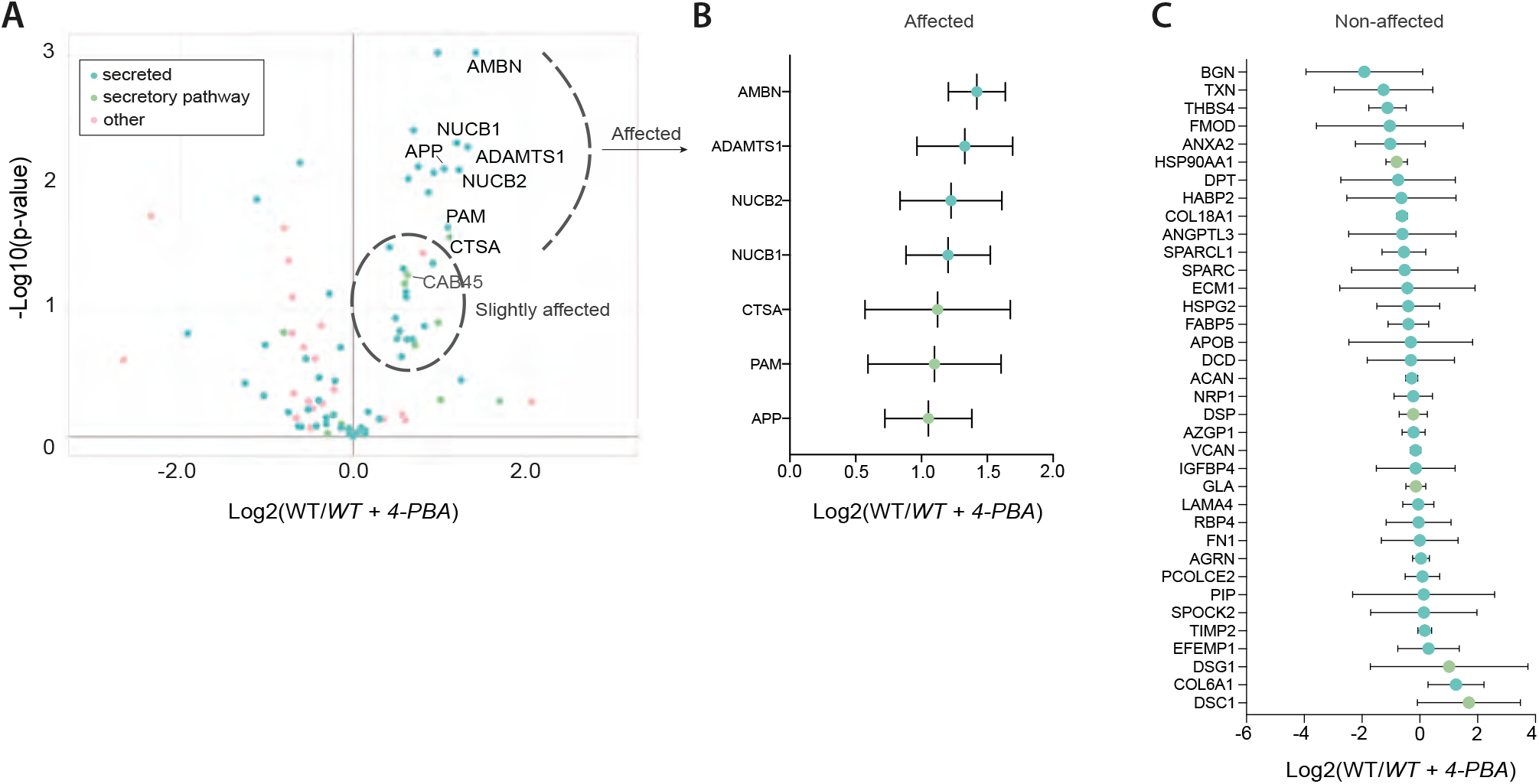
Proteomic analysis of secretomes from 4-PBA treated cells. **(A)** Volcano plot showing changes in protein secretion upon 4-PBA treatment in HEK-293 TREx cells. Proteins were considered affected if Log2 SILAC ratio was >1 and p<0.05 **(B)** Bar graph showing proteins with significantly decreased secretion upon 4-PBA treatment. **(C)** Bar graph showing proteins whose secretion was not significantly affected by 4-PBA treatment.

**Figure S8:**
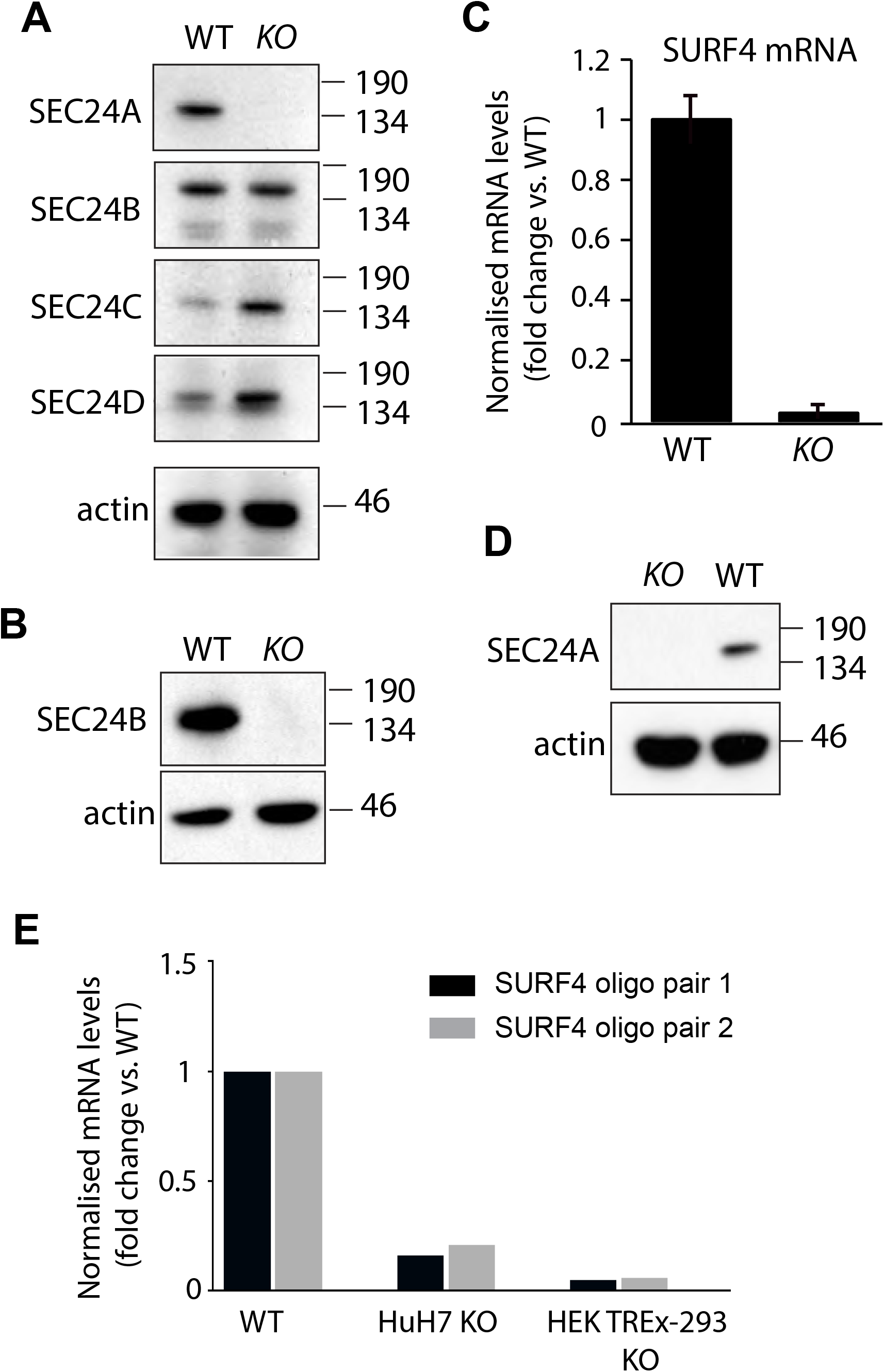
Validation of KO cell lines. **(A)** Western blot analysis of *SEC24A KO* CRISPR/Cas9 edited HEK-293 cells. Stability of Sec24 paralogs was analysed by immunoblotting cell lysates prepared from equal numbers of cells with specific antibodies (α-SEC24A, α-SEC24B, α-SEC24C and α-SEC24D) in WT and *SEC24A KO* cell line. α-actin was used as loading control. **(B)** Western blot analysis of *SEC24B KO* CRISPR/Cas9 edited HEK-293 cells. The presence of Sec24B was analysed by immunoblotting cell lysates prepared from equal numbers of cells with α-SEC24B antibodies. α-actin was used as loading control. **(C)** SURF4 mRNA expression in *SURF4 KO* CRISPR/Cas9 edited HEK-293 cells. Values are presented as mean ± SD of n = 3 independent experiments. **(D)** SURF4 mRNA expression in *SURF4 KO* CRISPR/Cas9 edited HuH7 cells. Values are presented as mean ± SD of n = 3 independent experiments.

